# Concatenation of segmented viral genomes for reassortment analysis

**DOI:** 10.1101/2024.02.26.582008

**Authors:** A. Ivanova, P. Volchkov, A. Deviatkin

**Affiliations:** Shemyakin-Ovchinnikov Institute of Bioorganic Chemistry, RAS (IBCh RAS), 117997 Moscow, Russia; Federal research center for innovator and emerging biomedical and pharmaceutical technologies, Russia, Moscow, 125315

## Abstract

Most reassortment identification methods are based on searching for phylogenetic discrepancies between phylogenetic trees for different segments. Other methods use pairwise genetic distances or compare the position of individual genome components in a tree relative to a reference component of the viral genome. However, such approaches are labour-intensive and hardly scalable.

Recent advances in the availability of viral sequencing technologies have led to the sequencing of large numbers of pathogen genomes, making manual processing of this large data difficult. At the same time, recombination analysis methods can process almost any number of sequences simultaneously. Such approaches are not suitable for the simultaneous analysis of multiple segments and are therefore not used to search for reassortment events. However, in the case of sequential concatenation of all segments, the methods of recombination analysis can be used to detect traces of reassortment events.

The code is available at https://github.com/melibrun/Concatenation-of-segmented-viral-genomes-for-reassortment-analysis. The service is implemented as a web application https://melibrun.shinyapps.io/viralsegmentconcatenator1/. It concatenates segmented viral genomes for reassortment analysis. The tool accepts files in GenBank format as input and generates a set of sequences in fasta format that are sequentially concatenated sequences of viral segments named in accordance with the “strain” field of the GenBank record annotation.

In order to use recombination search algorithms in the study of reassortment events, we have developed a method (Virus Segment Concatenator, VSC) to automatically concatenate the sequences of all segments of a virus into a single sequence.

The applicability of VSC for automated searches for reassortment events was demonstrated using CCHFV, an H5N5 subtype of influenza virus.

## Introduction

RNA viruses have different genome organisations. Some families are characterised by a segmented genome (e.g. Orthomyxoviridae), while others have a non-segmented genome (e.g. Rhabdovirus). At the same time, several members of the family *Chuviridae* have a segmented and a non-segmented genome (Li et al., 2015). Recently, new viruses with segmented genomes have been described from the *Flaviviridae* family. The diversity of viruses with segmented genomes is thus greater than previously assumed (Shi et al., 2016).

Reassortment is a segmental exchange between related viruses in coinfected cells (Lowen, 2018). This is a more common process than recombination. For example, in the 1950s, reassortment occurred between the neuraminidase of avian and human influenza A viruses (IAVs), leading to the emergence of ‘Asian’ influenza. In 2009, North American H3N2 and H1N2 swine viruses reassorted with Eurasian avian-like swine viruses and caused a pandemic (Neumann, Noda & Kawaoka, 2009).Software for finding reassortments usually requires the use of each segment. Many of these methods are based on phylogenetic mismatches between segments, so these differences can be highlighted and identified using tree-based approaches. Comparing differences between trees using tanglegrams or distance-based subtree prune and re-graft methods (Svinti, Cotton & McInerney, 2013; Varsani et al., 2018) provides deep insights into the analysis of reassortment events. Many of these methods are implemented in the tools, for example SMARTIE (Stochastic model for reassortment and transfer events) (Bloomquist & Suchard, 2010) and GIRaF (Graph-incompatibility-based Reassortment Finder) (Nagarajan & Kingsford, 2011). Other methods use pairwise genetic distances or compare where individual genome components are located in the tree in relation to the reference component of the viral genome (Varsani et al., 2018).

At the same time, there are several recognised methods for recombination analysis. For example, recombination detection program, RDP5 (Martin et al., 2021), Genetic Algorithm Recombination Detection, GARD (Kosakovsky Pond et al., 2006), pairwise distance divergence matrices (PDDM) (Vakulenko et al., 2021) make it possible to detect signs of recombination in the sequence alignment using different methods. It should be noted that such approaches are not able to analyse multiple segments simultaneously, indicating intersegmental phylogenetic incongruence. Technically, however, a variety of recombination analysis methods apply to reassortment analysis when all segments are sequentially concatenated. At the same time, recent advances in viral sequencing have resulted in a huge number of known pathogen genomes, making manual processing of this large data challenging. To this end, we have developed the Virus Segment Concatenator, VSC, (https://github.com/melibrun/Concatenation-of-segmented-viral-genomes-for-reassortment-analysis/tree/main). This software uses the Genbank file of the target taxon, selects viruses for which complete genomes have been sequenced, and concatenates the segments one by one. Virus Segment Concatenator (VSC) outputs putative complete viral genome sequences in fasta format, following the principle of “one virus -one sequence”.

## Materials & Methods

VSC concatenate segmented viral genomes for reassortment analysis. The tool accepts files in GenBank format as input. The default values are set for the influenza A virus. If the subject is another virus, should appropriate number of segments and range of segment lengths that we recognise as fully sequenced should be selected. As a result, the tool generates a set of sequences in fasta format that are sequentially concatenated sequences of viral segments named in accordance with the “strain” field of the GenBank record annotation. Additionally, the option to choose only protein-coding sequences has been added to the tool. In the case of the presence of several protein coding sequences the longest one is selected. Noteworthy all examples demonstrated in this study has been processed using this option.

All alignments were created using the default settings of the MAFFT online server (Katoh & Standley, 2013). Phylogenetic trees for the sequences were constructed using the neighbour-joining algorithm implemented in MEGA7(Kumar, Stecher & Tamura, 2016). The percentage of replicate trees in which the associated taxa clustered in the bootstrap test (1000 replicates) is shown next to the branches. The tree is drawn to scale, with branch lengths in the same units as the evolutionary distances used to infer the phylogenetic tree. The evolutionary distances were calculated using the Tajima-Nei method and are given in the unit of the number of base substitutions per site.

### H5N5 dataset preparation

Novel reassortant viruses have been described for the H5N5 influenza virus (Gu et al., 2011). To demonstrate the possibility of an automated search for reassortments, this subtype of influenza virus was selected. To uncover patterns of reassortment events in the H5N5 subtype, we downloaded all sequences available in Genbank as of 10 June 2023 that were annotated as belonging to the H5N5 subtype (n = 343). For this purpose, sequences were filtered that were annotated as belonging to txid465975, the taxonomy identification number of H5N5 subtype. In addition, these sequences were concatenated with VSC. In total, 40 viruses with known complete genomes were identified.

The concatenated sequences were analysed using RDP5 software and the approach proposed in (Vakulenko et al., 2021).

### CCHFV dataset preparation

In order to demonstrate the universality of the proposed approach, we chose a CCHFV whose genome consists of three segments. This virus is a representative of Bunyaviruses, that can cause diseases such as severe fever with thrombocytopenia syndrome, Rift Valley fever and Bwamba fever. Reassortment is thought to complement genetic drift as it is an important mechanism for the evolution of the bunyaviruses (Briese, Calisher & Higgs, 2013). In our previous research the abundant reassortment for the CCHFV has been demonstrated (Lukashev & Deviatkin, 2018).Here we have concatenated all available (as of 10 June 2023) complete genome sequences of CCHFV in GenBank using VSC. For this purpose, sequences were filtered that were annotated as belonging to txid3052518, the taxonomy identification number of *Orthonairovirus haemorrhagiae* (CCHFV).

In total, 179 viruses with known complete and near complete genomes were identified [. Sequence segment has been identified as “complete” or “near complete” if S segment has length 1500-1700 nucleotides, M segment - 5200-5400 nucleotides, L segment - 11900-12100 nucleotides.

### Data availability

The code has been deposited to https://github.com/melibrun/Concatenation-of-segmented-viral-genomes-for-reassortment-analysis [https://www.doi.org/10.5281/zenodo.10406114].

*H5N5* sequences used for the analysis are available at https://github.com/melibrun/Concatenation-of-segmented-viral-genomes-for-reassortment-analysis/blob/main/examples/Influenza%20A%20virus/H5N5.gb.

CCHFV sequences used for the analysis are available at https://github.com/melibrun/Concatenation-of-segmented-viral-genomes-for-reassortment-analysis/blob/main/examples/Crimean-Congo%20hemorrhagic%20fever%20orthonairovirus/Crimean-Congo%20hemorrhagic%20fever%20orthonairovirus.gb.

The code of the tool has been written in Python, available at https://github.com/melibrun/Concatenation-of-segmented-viral-genomes-for-reassortment-analysis. This code was implemented in the R shiny webserver as an online tool available at https://melibrun.shinyapps.io/combined_the_segments/.

Phylogenetic trees from this study are available at https://github.com/melibrun/Concatenation-of-segmented-viral-genomes-for-reassortment-analysis/blob/main/examples/Influenza%20A%20virus/PA.nwk; https://github.com/melibrun/Concatenation-of-segmented-viral-genomes-for-reassortment-analysis/blob/main/examples/Influenza%20A%20virus/PB1.nwk; https://github.com/melibrun/Concatenation-of-segmented-viral-genomes-for-reassortment-analysis/blob/main/examples/Crimean-Congo%20hemorrhagic%20fever%20orthonairovirus/S.nwk; https://github.com/melibrun/Concatenation-of-segmented-viral-genomes-for-reassortment-analysis/blob/main/examples/Crimean-Congo%20hemorrhagic%20fever%20orthonairovirus/M.nwk.

The RDP5 output for H5N5 is available at https://github.com/melibrun/Concatenation-of-segmented-viral-genomes-for-reassortment-analysis/blob/main/examples/Influenza%20A%20virus/H5N5.rdp5.

The RDP5 output for CCHFV is available at https://github.com/melibrun/Concatenation-of-segmented-viral-genomes-for-reassortment-analysis/blob/main/examples/Crimean-Congo%20hemorrhagic%20fever%20orthonairovirus/CCHFV.rdp5.

The sequences used for the reassortment analysis for the H5N5 dataset are available at https://github.com/melibrun/Concatenation-of-segmented-viral-genomes-for-reassortment-analysis/blob/main/examples/Influenza%20A%20virus/H5N5_event1.fas.

The sequences used for the reassortment analysis for the CCHFV dataset are available at https://github.com/melibrun/Concatenation-of-segmented-viral-genomes-for-reassortment-analysis/blob/main/examples/Crimean-Congo%20hemorrhagic%20fever%20orthonairovirus/cchfv_event1.fas.

### Description of the VSC

VSC enables the automatic concatenation of sequences written in GenBank or FASTA format. Please note that for correct analysis of FASTA sequences, the header should be written as follows: “>((strain name(subtype))_segment_segment-number” (e.g. “>(A/waterfowl/Korea/S005/2014(H5N5))_segment_3”). Sample file in FASTA format can be found at https://github.com/melibrun/Concatenation-of-segmented-viral-genomes-for-reassortment-analysis/blob/main/examples/Influenza%20A%20virus/H5N5_gisaid.fasta.

To visualise the separation between the segments, an option has also been added to add a certain number of “-” symbols.

If the name of the subtype is not specified in the GeneBank field “Organism”, the button “Please indicate whether the name of the subtype is specified in the ORGANISM field in the GenBank record” should not be activated. The name of the subtype should be written in the field “Please indicate the name of the taxon of interest”. Otherwise, if the name of the subtype is written in the GenBank field “Organism”, the button “Please indicate whether the name of the subtype is specified in the ORGANISM field in the GenBank record” should be activated and the field “Please indicate the name of the taxon of interest” should be empty.

The default settings of the application are set for the influenza A viruses whose genome consists of eight segments of the specified length. If the virus of interest has a different number of segments with a different length, this should be specified in the appropriate fields.

The “Permissible difference” fields specify the range of segment length that will be treated as a complete segment sequence. For example, if “Permissible difference” is 200 and the length of segment 1 is 2300, sequences with a length between 2100 and 2500 will be treated as a complete segment length.

The use of the annotated GeneBank file enables the use of additional options. For example, only protein-coding regions can be concatenated if the corresponding button has been selected.

## Results

### Avian Influenza (H5N5) Viruses

VSC has been used for the dataset of H5N5 IAVs. The resulting dataset contained 40 concatenated genomes of these viruses. According to RDP5, six reassortment events were validated by at least six of the seven recombination identification algorithms.

A comprehensive snapshot of the reassortment within this influenza subtype was represented by pairwise distance divergence matrices (PDDM) (Vakulenko et al., 2021). In this plot, the regions between which reassortment has occurred between the most divergent viruses are colour-coded (Figure 1). According to this result, the most distant H5N5 viruses preferentially exchange segments 4 (HA), 6 (NA) and 8 (NS). However, warm colours are also found in the intersection of other segments. At the same time, cold colours are typical for the comparison of intrasegment sequences, indicating the absence of recombination traces.

**Figure 1.**
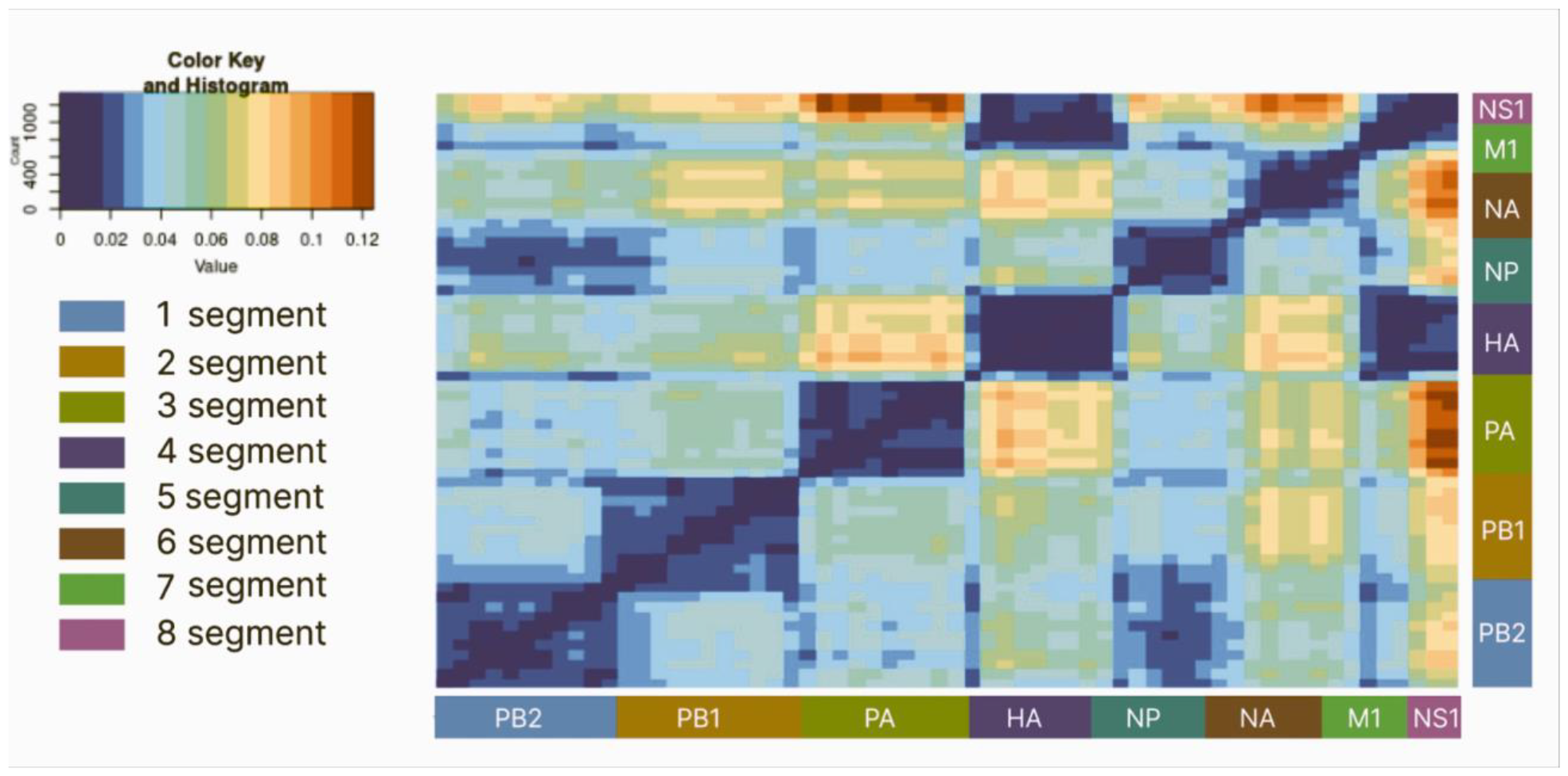
Pairwise distance divergence matrices (PDDM) for H5N5 influenza viruses (Window = 500, Step = 200). The PDDMs were constructed based on the deviations of the dots in the pairwise distance correspondence plots (PDCPs) of the relevant genomic regions. The colour gradient shows the root-mean-square error (RMSE) values in the PDCPs.

To reveal concrete reassortant viruses we have built pairwise distance correspondence plots (PDCP) between 2nd (2281-4554 nt in alignment https://github.(Vakulenko et al., 2021)<u>com/melibrun/Concatenation-of-segmented-viral-genomes-for-reassortment-analysis/blob/main/examples/Influenza%20A%20virus/H5N5_alig.fasta</u>)and 3rd segment (4558-6708 nt in alignment https://github.com/melibrun/Concatenation-of-segmented-viral-genomes-for-reassortment-analysis/blob/main/examples/Influenza%20A%20virus/H5N5_alig.fasta) (Figure 2B). For example, 2nd segment, “PB1”, of A/duck/Eastern_China/031/2009 and A/duck/EasternChina/008/2008 shares 99,6 % of identical nucleotides, whereas 3rd segment, “PA”, shares 94,3 % of identical nucleotides.

**Figure 2.**
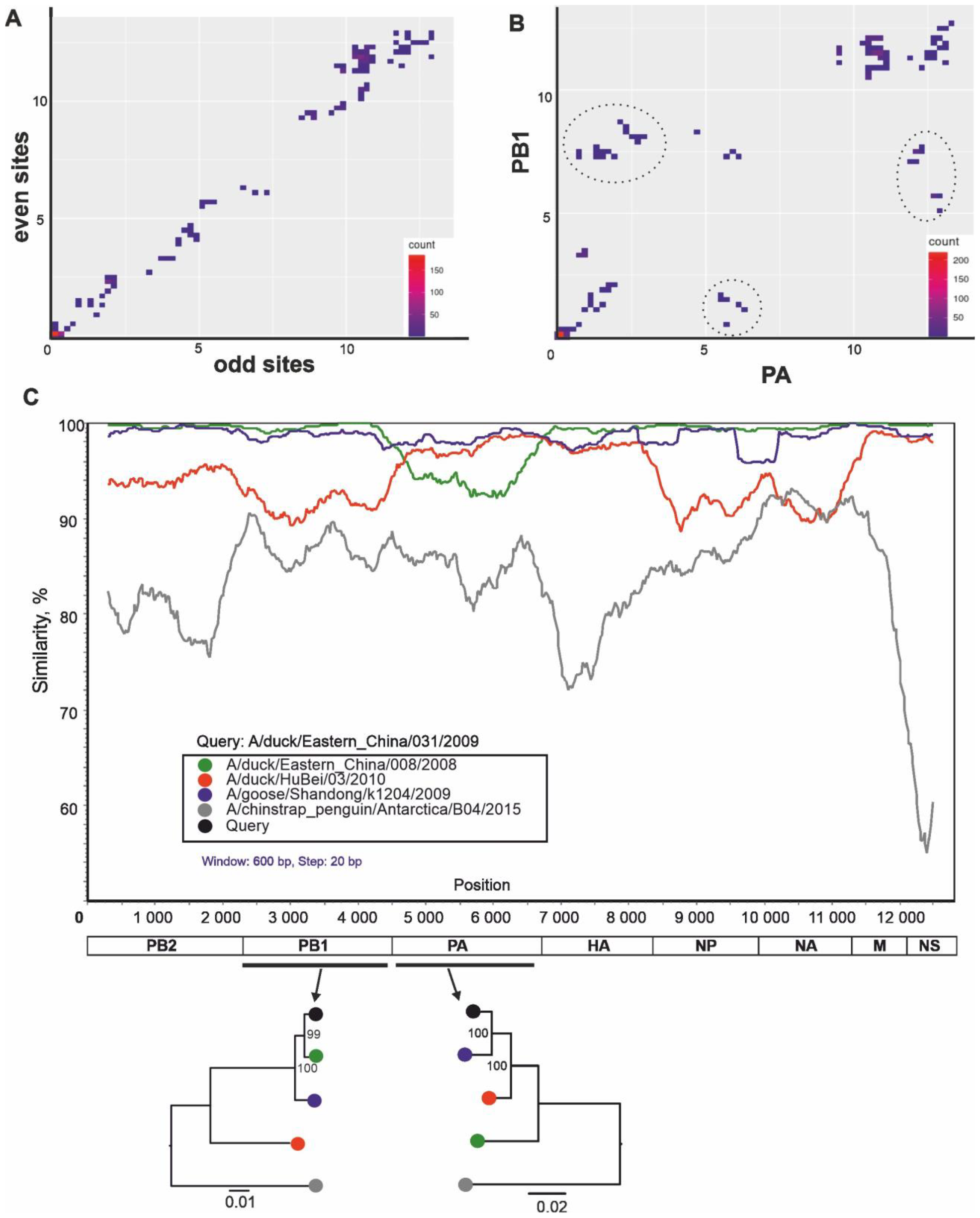
Pairwise nucleotide distance comparison plots (PDCP) demonstrate phylogenetic incongruence between selected genetic regions of H5N5 influenza viruses. Each dot indicate a pair of raw nucleotide distances between the same pair of genomes in two genomic regions (axis labels). A. PDCP of even for concatenated 2nd (PB1) and 3rd (PA) segments for which distances were calculated for even and odd sites (negative control); B. PDCP between PB1 and PA segments of H5N5 influenza viruses. The dashed circles indicate virus pairs where the difference in genetic distances was evident; C. Similarity plot demonstrate complex evolutionary history patterns in H5N5 influenza viruses. The similarity scan was performed using alignments of five nucleotide sequences indicated in the legend with a window/step size of 600/20 nt.

This plot demonstrated that these segments of many viruses has divergent origin. To demonstrate negative control, we concatenated the PB1 and PA segments under investigation and calculated the pairwise distances for even (y-axis) and odd (x-axis) positions (Figure 2A). As a result, as expected, a linear trend was observed -the more substitutions occurred at the even positions of the concatenation, the more substitutions occurred at the odd positions of the concatenation.

Some of such reassortment viruses has been revealed by similarity plots and phylogenetic trees (Figure 2C).

### Crimean-Congo hemorrhagic fever orthonairovirus

The concatenated sequences of CCHFV were analysed using RDP5 software (Martin et al., 2021) and the approach proposed in (Vakulenko et al., 2021). In total, 33 reassortment events have been validated by at least six of the seven recombination identification algorithms. A comprehensive snapshot of reassortment within CCHFV was represented by pairwise distance divergence matrices (PDDM) (Vakulenko et al., 2021) (Figure 3). In this plot, the regions between which reassortment has occurred between the most divergent viruses are colour-coded (Figure 3). According to this result, the most distant CCHFV preferentially exchange S and L segments and M segment. It should be noted, that the first fragment of M segment (nearly first 1000 nucleotides) and last fragment of M segment have diverse origin.

**Figure 3.**
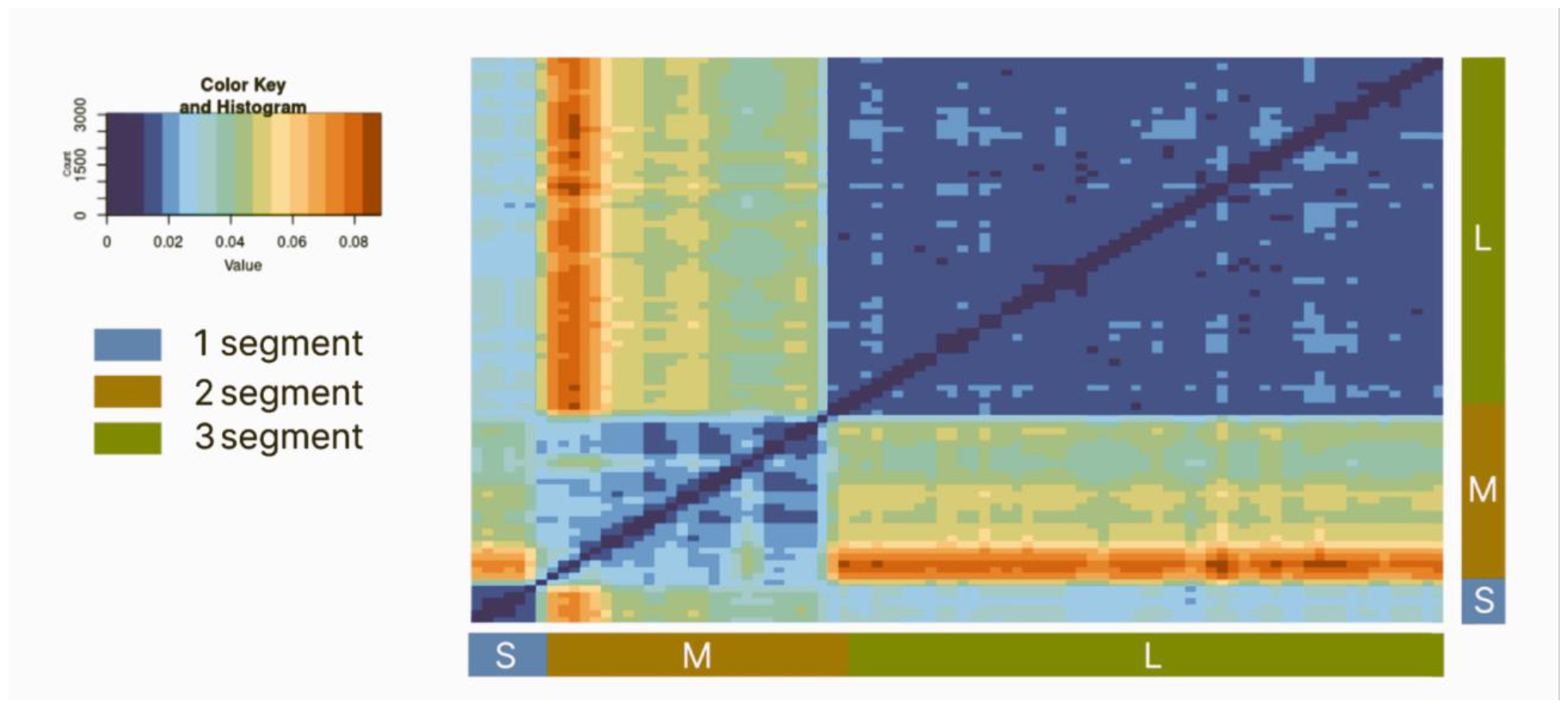
Pairwise distance divergence matrices (PDDM) for CCHFV (Window = 500, Step = 200). The PDDMs were constructed based on the deviations of the dots in the PDCPs of the relevant genomic regions. The colour gradient shows the root-mean-square error (RMSE) values in the PDCPs.

To reveal concrete reassortant viruses we have built pairwise distance correspondence plots (PDCP) between S segment (1-1470 nt in alignment https://github.com/melibrun/Concatenation-of-segmented-viral-genomes-for-reassortment-analysis/blob/main/examples/Crimean-Congo%20hemorrhagic%20fever%20orthonairovirus/crim_aligned.fasta) and first part of M segment (1471-2500 nt in alignment https://github.com/melibrun/Concatenation-of-segmented-viral-genomes-for-reassortment-analysis/blob/main/examples/Crimean-Congo%20hemorrhagic%20fever%20orthonairovirus/crim_aligned.fasta) (Figure 4B). For example, S segments of 2014-C-K14-1C and MCL-19-T-1916 viruses shares 88,9% of identical nucleotides, whereas first part of M segments shares 51,7% identical nucleotides. This plot demonstrated that these segments of many virus pairs had divergent origin. Some of such reassortment viruses has been revealed by similarity plots (Figure 4C) and phylogenetic trees.

**Figure 4.**
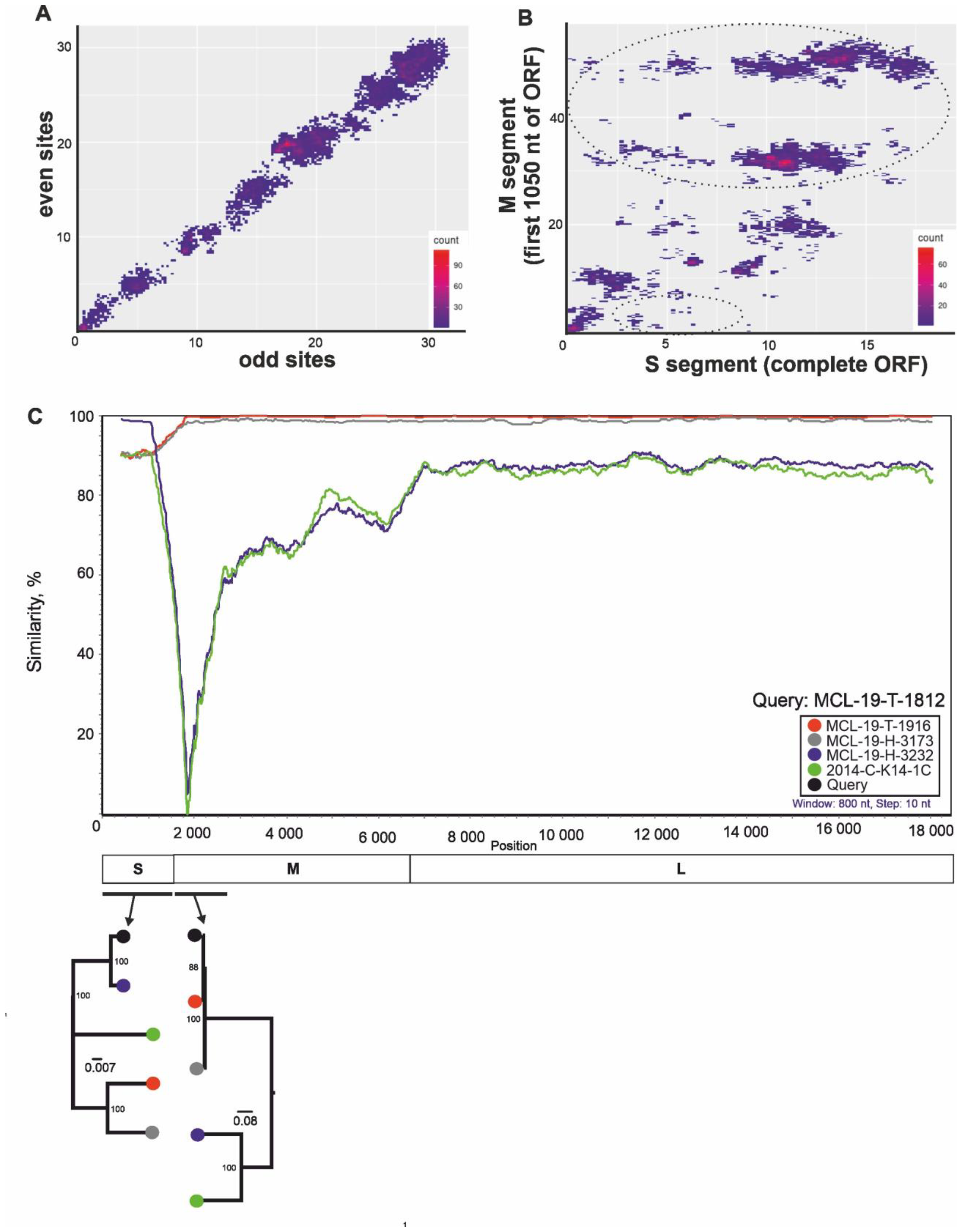
Pairwise nucleotide distance comparison plots (PDCP) demonstrate phylogenetic incongruence between selected genetic regions of CCHFV. Each dot indicate a pair of raw nucleotide distances between the same pair of genomes in two genomic regions (axis labels). A. PDCP of even for concatenated S and first part of M segments, for which distances were calculated for even and odd sites (negative control); B. PDCP between S and first part of M segments of CCHFV. The dashed circles indicate virus pairs where the difference in genetic distances was evident; C. Similarity plot demonstrate complex evolutionary history patterns in CCHFV. Similarity scan was performed using alignments of six nucleotide sequences indicated in the legend with window/step size 800/10 nt.

## Discussion

In the last ten years, advances in sequencing technologies have significantly increased the number of known sequences in publicly available databases. For example, in January 2001 there were 3,226 sequences of IAVs, in January 2010 - 125,910 sequences of IAVs, in January 2023 - 871,664 sequences in the GenBank database. In itself, a large amount of information is an important contribution to the overall knowledge about the diversity of viruses. However, this wealth of information is not without its intricacies. For example, newly sequenced viruses should be compared with a complete data set of related viruses in order to obtain comprehensive rather than fragmentary results.

Recombination and reassortment can lead to the emergence of a virus whose genomic fragments have different origins (Varsani et al., 2018). Currently, recombination can be automatically detected for a huge data set by various approaches (Martin et al., 2015; Vakulenko et al., 2021). At the same time, to our knowledge, there is no robust method for automatically searching for reassortment events. To solve this problem, we propose to use VSC as the first step of a comprehensive analysis of reassortment in a dataset of virtually any size.

It should be noted that the commercially available Geneious software (“https://www.geneious.com/“) provides an option to concatenate the sequences. However, this requires the addition of the standardized names to each sequence, which precludes the use of this software for the automatic detection of reassortment events. In contrast, VSC allows concatenation of the viral segments with sequences in GenBank raw format.

Similar approach of segment concatenation has been applied for the reassortment analysis in Banana bunchy top virus (Stainton et al., 2012, 2015). At the same time VSC allows to conduct such analysis automatically without any need in additional data parsing.

In this study, we demonstrated the ability of reassortment screening based on segmental concatenation using a dataset that contained previously characterised reassortant H5N5 viruses(Gu et al., 2011) (positive control) (Figures 1,2). Our results were consistent with previous findings. To verify whether such an approach is capable of finding yet unknown reassortment events, we assembled all available sequences of CCHFV and analysed them with PDDM and PDCP (Figures 3 and 4). As a result, the novel reassortant virus MCL-19-T-1812 was uncovered, which had not been detected before (Sahay et al., 2020). In other words, the proposed approach makes it possible to find reassortant viruses based on the automated analysis steps. It should be noted that VSC is a tool that automatically concatenates sequential genome segments of viruses with segmented genomes. This is a preparatory and necessary step for the ability to apply the recombination search algorithms for reassortment analysis.

## Conclusions

In recent years, the number of known viral sequences of epidemiological importance has increased considerably. At the same time, reassortment plays an important role in the emergence of fundamentally new variants of viruses with segmented genomes. However, there are currently no approaches that allow automated processing of large data sets with genomic sequences of different viral segments without any preprocessing. The current software solution, which was demonstrated at two different objects, offers a new approach for automated reassortment searches.

